# HextractoR: an R package for automatic extraction of hairpins from genome-wide data

**DOI:** 10.1101/2020.10.09.333898

**Authors:** Cristian Yones, Natalia Macchiaroli, Laura Kamenetzky, Georgina Stegmayer, Diego Milone

## Abstract

Extracting stem-loop sequences (hairpins) from genome-wide data is very important nowadays for some data mining tasks in bioinformatics. The genome preprocessing is very important because it has a strong influence on the later steps and the final results. For example, for novel miRNA prediction, all well-known hairpins must be properly located. Although there are some scripts that can be adapted and put together to achieve this task, they are outdated, none of them guarantees finding correspondence to well-known structures in the genome under analysis, and they do not take advantage of the latest advances in secondary structure prediction. We present here an R package for automatic extraction of hairpins from genome-wide data (HextractorR). HextractoR makes an exhaustive and smart analysis of the genome in order to obtain a very good set of short sequences for further processing. Moreover, genomes can be processed in parallel and with low memory requirements. Results obtained showed that HextractoR has effectively outperformed other methods.

HextractoR it is freely available at [CRAN](https://cran.r-project.org/web/packages/HextractoR/index.html) and [Sourceforge](https://sourceforge.net/projects/sourcesinc/files/hextractor/).

## 1 Introduction

Extracting stem-loop sequences (hairpins) from genome-wide data is very important for some data mining tasks in bioinformatics such as the computational prediction of pre-microRNAs (pre-miRNAs) with machine learning. In most works (Demirci *et al*., 2017; Stegmayer *et al*., 2018) the datasets used to test the prediction methods are manually built, using several interconnected tools. This has the obvious disadvantage of requiring a not negligible manual amount of work. Moreover, the process has a great impact in the prediction task afterwards. If some stem-loops are not correctly identified and extracted from the genome, the prediction method will not be able to detect the corresponding sequence. If they are detected but incorrectly trimmed (longer or shorter than a corresponding pre-miRNA) the features extracted from these sequences can vary a lot, making machine learning prediction methods to generate incorrect predictions. There are other tools to extract hairpins (Yang and Li, 2011; Friedländer *et al*., 2008) but they use RNAseq data and they are not designed to extract all hairpins, only the ones that match with a significant number of reads. Finally, since in most works this first stage of stem-loop extraction is performed manually by combining several tools, it is very difficult, or even impossible, to reproduce the results. For this reason the experiments of most published pre-miRNA prediction methods cannot be reproduced accurately., and therefore the users of those tools cannot obtain the same prediction rates published. With HextractoR, besides providing a unique tool to simplify this stage of pre-miRNA prediction, a standardized way to perform this important preprocessing task is proposed. After hairpins extraction, the miRNAfe tool (Yones *et al*., 2015) can be used, which combines all the features previously described for pre-miRNAs in a single tool. It is actually being used in most recent prediction models (Yones *et al*., 2017; Acar *et al*., 2018). Furthermore, it has been already used successfully for building 6 public genome-wide datasets Bugnon *et al*. (2019). HextractoR helps to standardize and simplify the stem-loop extraction stage, making future prediction methods easier to use and their experiments fully reproducible.

## 2 HextractoR pipeline

HextractoR predicts the secondary structure of several overlapped segments, with a longer length than the mean length of the sequences of interest for the species under processing, ensuring that no one is lost nor inappropriately cut. The length of the cutting window can be configured to define the maximum size that the stems found will have (smaller stems will also be found). Optionally, FASTA files containing known sequences can be loaded and used to split the output stem-loops into several FASTA output files, according to their type. If no filter file is provided HextractoR generates just one FASTA file with all hairpins. If two filter files are provided, for example, with well-known pre-miRNAs and other with known non-miRNA sequences, HextractoR generates three FASTA files: one for each filtering file passed to HextractoR with the sequences that match according to BLAST (Altschul *et al*., 1990) with known sequences; and another one with the stem loops that did not match. The processing steps of HextractoR (see Figure 1) are explained in detail in the next sections.

**Figure 1:**
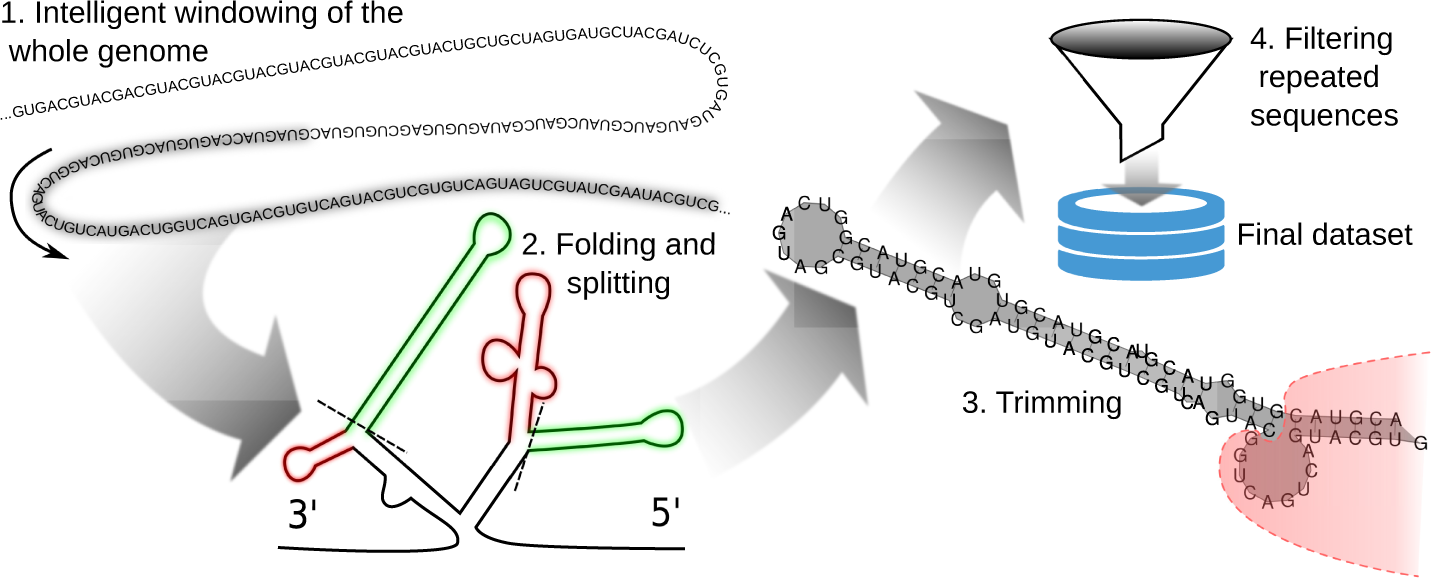
The HextractoR pipeline.

The input parameters of HextractoR are:

- Minimum number of valid nucleotides required to process a sequence: to avoid the processing of sequences too shorts or with many non identified nucleotides (the Ns).
- Length and step of the cutting window. The step is the number of positions for sliding the window.
- Minimum hairpin length and minimum number of base-pairs.
- A boolean indicating if trimming is required. If not, the whole stem-loops are returned.
- The e-value and identity percentage used when comparing the known sequences with the stem-loops extracted.
- The number of threads used for parallel execution.

Optionally, one or several filtering fasta files containing well-known sequences can be loaded and used to split the output stem-loops into several fasta files, according to their type. As output, HextractoR returns several fasta files: one for each filtering file passed to HextractoR with the sequences that match Altschul *et al*. (1990) with wellknown sequences; and another one called “unlabeled” for the stem loops that could not be matched.

The processing steps of HextractoR are (see Figure1):

### 2.1 Intelligent windowing of the whole genome

HextractoR starts by cutting the complete genome into overlapping windows of a large length (∼ 500 nt). The window must be long enough in order to correctly capture a complete hairpin, but also to take into account the neighborhood of any possible hairpin when estimating the secondary structure. This is very important since the results of estimating a secondary structure can be greatly affected by the neighborhood of the sequences.

### 2.2 Folding and splitting

The second step involves correctly predicting the secondary structure of the sequences obtained in the previous windowing step, when folding. To do this, the minimum free energy algorithm (Zuker and Stiegler, 1981) of RNAfold is used. This algorithm uses dynamic programming for finding the secondary structure that minimizes the energy released. Since the windows used are relatively long, the structures found usually have multiple loops. Therefore, they have to be split into several hairpin-type structures, as can be seen in Figure 1. Those hairpins that do not exceed a minimum length and level of pairing are eliminated.

### 2.3 Trimming

Since sequence windows with a length greater than the average of the sequence of interest are used, generally large secondary structures are obtained. For this reason, it is important to analyze each sequence in order to detect if trimming is necessary. Certain heuristics can be used to obtain sequences with lengths and stability properties similar to those of a well-known pre-miRNA. These rules optimize the Minimum Free Energy normalized by the sequence length (NMFE). Although optimum cutting points can be found by re-estimating the secondary structure for all possible cuts, a set of rules provide more flexibility and accelerate the process. First, the sequence must exceed a minimum length, pre-defined according to the species under study. In this way, it can be ensured that the secondary structure has sufficient length to be a pre-miRNA of the species under analysis. Secondly, the cuts are made in the first unpaired nucleotide of an internal loop or bulge of the secondary structure (starting from the main loop). To choose the bulge/loop where to cut, a score is assigned to each imperfection as

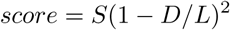

where *S* is the imperfection length, *D* is the distance to the main loop and *L* is the stem length (from the main loop to the last paired nucleotide). If the imperfections in the secondary structure are large, it is likely that cutting the sequence at those points will result in a structure with lower NMFE. The score favors larger imperfections. Moreover, the smaller the length of the sequence (independently of the pairing), the higher the NMFE. Therefore, a loop/bulge closer to the main loop is preferred.

### 2.4 Filtering repeated sequences

Repeated sequences are eliminated to avoid extra computational cost. These repeated sequences might also disturb the results of the prediction algorithms. Repetitions may appear due to the overlapping in windowing. These repeated sequences appear consecutively and they are almost identical sequences. To eliminate them, a comparison between each sequence and the last extracted sequence is made. If one of the sequences contains the other one, the shorter one can be discarded.

## 3 Complete genomes results

To test the R package, several genomes were processed and the results were compared to those obtained from using some mirCheck scripts (Jones-Rhoades and Bartel, 2004). There are many miRNA prediction methods available, but none of them provide the tools needed to extract stem-loops from a whole-genome. The mirCheck scripts were the only tool that we have found to extract stem-loop sequences from genome-wide data. After this extraction stage, a miRNA prediction method would be used to discard non miRNA hairpins. Some of these methods take advantage of all the stem-loop sequences (even the unlabelled) to improve the results, as those based on semi-supervised learning Yones *et al*. (2017). Moreover, to correctly assess the performance of miRNA prediction methods, it is important to have the widest variety of negative examples. Therefore, it is important to point out that the objective of HExtractoR is finding the largest possible number of hairpins in the genome (and not to classify or predict whether a given sequence is a miRNA). For this reason, some measures, such as the precision, are not relevant when testing HextractoR, as all stem-loop are considered worthwhile. Even more important than the number of stem-loops, is that all known pre-miRNAs appear among the extracted stem-loops. This results are shown in Table 1 for six species, comparing the number of hairpin found by each method. The number of pre-miRNAs available (according to miRBase v21(Griffiths-Jones *et al*., 2006)) for all species is reported in the fourth column of the table. The proportion of pre-miRNAs found by both methods is shown in the last two columns. It can be clearly seen that, in all cases, the performance of HextractoR is superior to mirCheck.

**Table 1:** Number of stem loops and pre-miRNA found with each tool.

| Species | Extracted hairpins |  | pre-miRNAs<br>in miRBase | pre-miRNAs found |  |
| --- | --- | --- | --- | --- | --- |
|  | mirCheck | HExtractoR |  | mirCheck | HExtractoR |
| *A. gambiae* | 1,410,532 | **4,276,543** | 66 | 92.42 % | **100.00 %** |
| *C. elegans* | 875,588 | **1,739,124** | 250 | 90.00 % | **99.60 %** |
| *D. rerio* | 11,028,128 | **23,214,338** | 346 | 93.06 % | **99.42 %** |
| *E. multilocularis* | 509,530 | **1,898,911** | 22 | 81.81 % | **100.00 %** |
| *A. thaliana* | 874,320 | **1,357,455** | 325 | 91.69 % | **94.77 %** |
| *H. sapiens* | 18,654,426 | **48,206,494** | 1881 | 85.38 % | **97.45 %** |

## 4 Conclusions

We have developed a simple and integrated tool, the R package HextractoR, that automatically extracts and folds all possible hairpin sequences from genome-wide data. The genomes can be processed in parallel and with low memory requirements since it can automatically split large multi-FASTA files. The proposed computational method, takes advantage of the latest developments in secondary structure prediction. Results obtained showed that HextractoR has effectively outperformed the state-of-the-art.

## References

Acar, İ., Saçar Demirci, M., Groß, U., and Allmer, J. (2018). The expressed microRNA-mRNA interactions of toxoplasma gondii. Frontiers in microbiology, 8, 2630.

Altschul, S., Gish, W., Miller, W., Myers, E., and Lipman, D. (1990). Basic local alignment search tool. Journal of molecular biology, 215(3), 403–410.

Bugnon, L., Yones, C., Raad, J., Milone, D., and Stegmayer, G. (2019). Genome-wide hairpins datasets of animals and plants for novel mirna prediction. Data in Brief, page 104209.

Demirci, M., Baumbach, J., and Allmer, J. (2017). On the performance of pre-microRNA detection algorithms. Nature communications, 8(1), 330.

Friedländer, M., Chen, W., Adamidi, C., Maaskola, J., Einspanier, R., Knespel, S., and Rajewsky, N. (2008). Discovering micrornas from deep sequencing data using mirdeep. Nature biotechnology, 26(4), 407.

Griffiths-Jones, S., Grocock, R., Van Dongen, S., Bateman, A., and Enright, A. (2006). mirbase: microRNA sequences, targets and gene nomenclature. Nucleic acids research, 34(suppl 1), D140–D144.

Jones-Rhoades, M. W. and Bartel, D. P. (2004). Computational identification of plant micrornas and their targets, including a stress-induced mirna. Molecular cell, 14(6), 787–799.

Stegmayer, G., Di Persia, L., Rubiolo, M., Gerard, M., Pividori, M., Yones, C., Bugnon, L., Rodriguez, T., Raad, J., and Milone, D. (2018). Predicting novel microrna: a comprehensive comparison of machine learning approaches. Briefings in bioinformatics.

Yang, X. and Li, L. (2011). mirdeep-p: a computational tool for analyzing the microrna transcriptome in plants. Bioinformatics, 27(18), 2614–2615.

Yones, C., Stegmayer, G., Kamenetzky, L., and Milone, D. (2015). mirnafe: a comprehensive tool for feature extraction in microRNA prediction. Biosystems, 138, 1–5.

Yones, C., Stegmayer, G., and Milone, D. (2017). Genome-wide pre-mirna discovery from few labeled examples. Bioinformatics, 34(4), 541–549.

Zuker, M. and Stiegler, p. (1981). Optimal computer folding of large rna sequences using thermodynamics and auxiliary information. Nucleic acids research, 9(1), 133–148.

